# Representational capacity shapes both the “whether” and “how” of social learning

**DOI:** 10.64898/2026.03.07.710273

**Authors:** Max Taylor-Davies

## Abstract

Social learning is widely understood as offering a mechanism to mitigate the costs and risks of individual trial-and-error exploration. This cost-avoidance account implies a framing of social learning as a resource-rational adaptation, which should be most beneficial to populations with limited capacity to learn asocially. But in cases of peer-to-peer transmission, that same limited capacity may hinder the reliability of social information—rendering it less, not more, useful. So how do these conflicting intuitions resolve? Across a series of simulation experiments, we find evidence for an ‘inverse-U’ relationship where social information use emerges most strongly in populations with moderate capacity relative to the complexity of their environment. Furthermore, we demonstrate a co-evolutionary transition in social learning strategies: as population capacity rises, selection pressure shifts from favouring a success bias to a conformist approach. Our findings suggest that constraints on information-processing play an important role in the emergence of culture, determining not just if a species learns socially, but also how.

All code for this project is available at https://github.com/maxtaylordavies/capacity-and-social-learning

**Author summary:** Many animals learn not only from their own trial-and-error exploration, but also by observing others. This “social learning” can be extremely useful, allowing young or inexperienced individuals to avoid the dangers of interacting with the environment themselves. This may be especially true for animals that are less able to learn by themselves due to limited attention or memory. But if these same limits affect those animals’ peers, the social information available will become less reliable, making social learning less, not more, useful. To reconcile these two opposing factors, we used computer simulations of agents learning which unfamiliar mushrooms were edible. We varied how much information agents could observe about each mushroom, and whether they learned alone or from others. We found that the benefits of social learning were strongest for neither the most nor least capable agents, but rather in between, where others were reliable enough to help but individual learning was still difficult. We also found that the same intervention influenced the best “strategy” for choosing whom to learn from: when agents could observe little information, it was best to copy the single most successful individual; when they could observe more information, it was best to follow the majority. July 1, 2026 1/18

## Introduction

Why is social learning so taxonomically widespread [1–4]? What makes learning from others so beneficial, when most animals are perfectly capable of learning by themselves? The prevailing answer to this question goes back to Boyd & Richerson [5] and holds that asocial learning (via trial-and-error exploration) imposes certain costs; either through the baseline metabolic requirements of the exploration process itself, or more dramatically by exposing the learner to risks of serious harm. By leveraging the experience of their conspecifics, and thus reducing the amount of asocial exploration necessary to acquire adaptive behaviours, the social learner mitigates these costs to eke out a fitness advantage [6–8]. Take the example of mushroom foraging [9], where a novice must learn to consume safe species while avoiding poisonous ones. A trial-and-error approach, while not impossible, might kill an unlucky learner before they can reliably distinguish the two. More prudent would be to follow the cues given by an experienced forager, who (perhaps by learning from someone else in turn) already possess the relevant environmental knowledge [10–12].

Considering the general case where social information may be noisier or less reliable than direct first-hand experience, this cost-avoidance account bears a structural resemblance to the picture at the heart of resource-rational analysis—which hold that evolution shapes cognition towards strategies that trade off optimally between the rewards they produce and the costs involved in executing them [13–18]. This has led some to characterise social learning as being itself a resource-rational adaptation [19–21], with the implication that it should be most beneficial to agents with limited capacity to learn ‘by themselves’.

A large body of previous theoretical work has identified various factors that influence the adaptiveness of social learning. Much of this work has focused on the effect of environmental variability, finding for example that social learning is favoured when the rate of change is low enough that information does not become obsolete too quickly, but not so low that no learning is required [5, 22–25]. Other studies have investigated the impact of task difficulty (typically via the size of the search space or the quality of available asocial evidence), demonstrating consistently that social learning is more strongly favoured in harder task environments [26, 27]. Less attention has been paid to how properties of the learner themselves (rather than of the environment) affect the utility of learning from others, with consideration of this angle limited mostly to factors such as agents’ strategies for choosing whom to learn from or when to favour social over asocial learning, or the network structure by which they are connected [6, 28–30]. No prior work has yet given explicit consideration to the impact of limits on information-processing capacity. At first glance, it might be argued that this is just task difficulty under another name, and so does not merit separate study: as we decrease an agent’s ability to process information and represent their environment, we increase the effective difficulty of the asocial learning problem they face—rendering social learning more attractive. But unlike most instances of resource-rational analysis, where it can make sense to consider the computational bounds of a single agent in isolation, the picture here is complicated by the fact that any realistic treatment of social learning must assume that the agents *providing* the social information are constrained in the same manner; any such capacity limit thus impacts the system in two distinct ways. This produces conflicting intuitions: decreasing capacity makes the asocial learning problem harder, which should favour social learning; at the same time, it reduces the quality of social information, which should favour *asocial* learning.

In this paper, we attempt to untangle the outcome of these two effects through a series of computational experiments. Using a contextual bandit problem modelled on the mushroom foraging setting, we investigate how the fitness advantage conferred by social learning varies as a function of agents’ capacity to represent decision-relevant features, in environments of varying complexity. Our results suggest that the benefit of social learning neither increases nor decreases monotonically with capacity, but instead follows an ‘inverse-U’ relationship—where social information use emerges most strongly in populations of intermediate capacity relative to the complexity of their environment. Finally, we also investigate the relative performance of two social learning *strategies*, finding that increasing capacity produces a systematic transition from success bias to conformity.

## Methods

### Shared bandit problem

We model the foraging problem using a deterministic version of the *contextual bandit* paradigm [31], faced in parallel by each of a population of different agents. In a regular non-contextual bandit problem, an agent proceeds through a sequence of trials, at each of which they choose one of a number of ‘arms’ to pull, gaining some amount of reward as a result. The agent’s task is to maximise cumulative payoff by estimating the reward distribution associated with each arm. In a contextual bandit, the only difference is that the agent additionally receives a piece of ‘context’ *x*_*t*_ at the start of each trial, the values of which correspond in some way to different settings of arm-reward probabilities. The agent must therefore learn a mapping from context value to optimal arm choice (e.g. “when *x*_*t*_ = 0 pull arm 1, when *x*_*t*_ = 1 pull arm 2”).

In our task, each context value *x*_*t*_ corresponds to the distinguishing features of a particular mushroom species. Specifically, we say that each species is characterised by a vector of *n* binary features s *x* ∈ *X* _*n*_ = {™1, +1} ^*n*^ (i.e. an *n*-feature environment contains 2^*n*^ different species). These features can be interpreted as referring to things like the cap colour of the mushroom, whether it has spots, its size, etc. A deterministic function *f*_*θ*_ maps this feature space to a binary label of +1 (edible) or ™1 (poisonous):

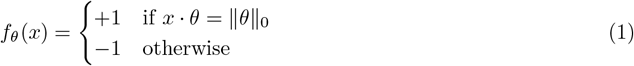

where∥·∥ _0_ returns the number of nonzero elements in its argument, and *θ*∈ Θ_*n*_ = {™1, 0, +1} ^*n*^ corresponds to a logical rule over some subset of context features. The allowance of zeros means that rules can ‘ignore’ certain features; for example, *θ* = +1 0 ™1 means that a species *x* must have *x*_1_ = +1, *x*_3_ = ™1 and either value of *x*_2_ in order to be classified as edible (see Fig 1A for an illustration of this example). We note also that *θ* is fixed for each instantiation of the task environment, so that a given population only ever experiences a single rule; all our results will be reported for simulation runs aggregated over many rules sampled uniformly from Θ_*n*_.

**Fig 1.**
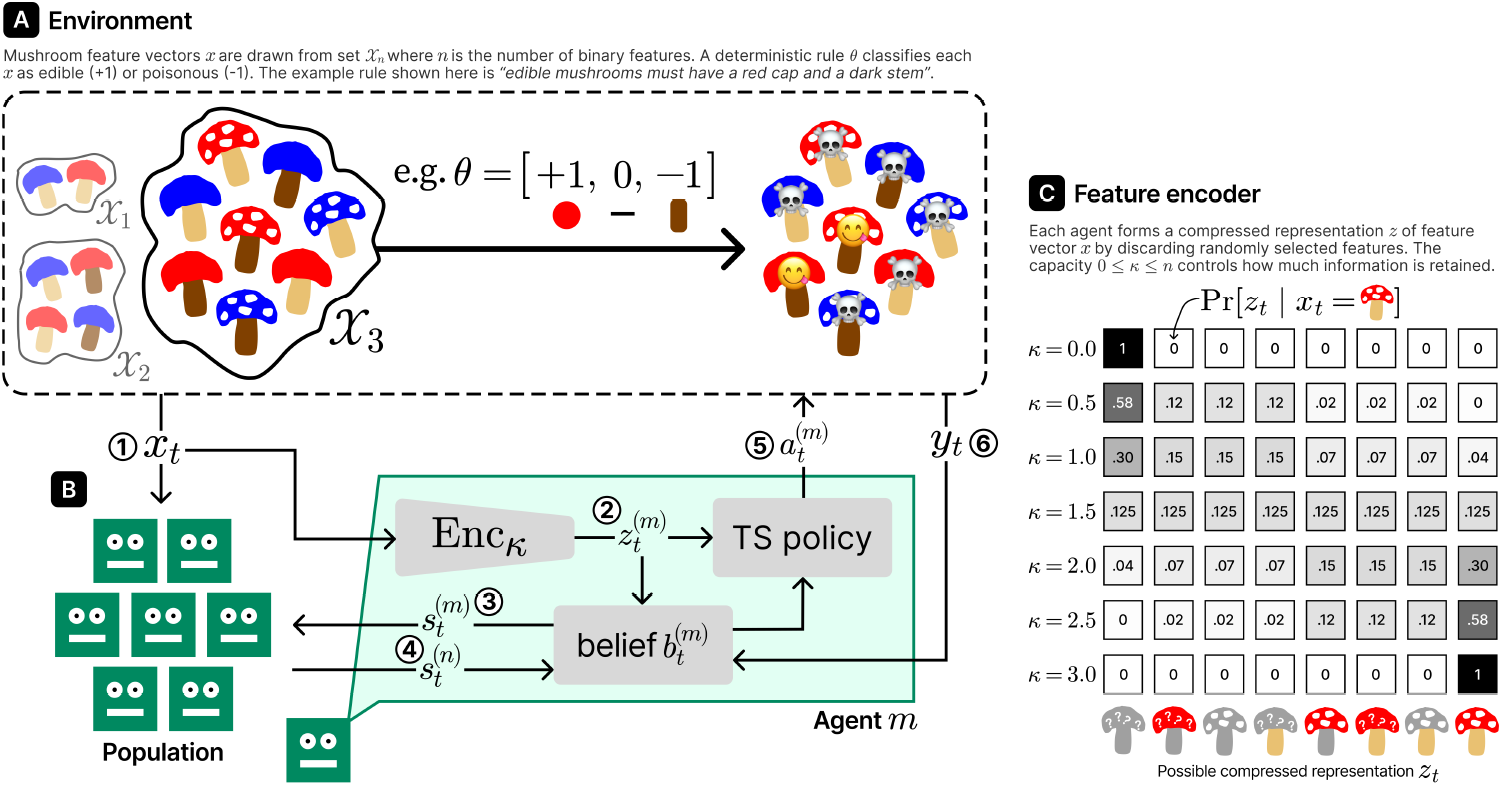
A high-level diagram of the bandit task and agent model used in our simulations. **(A)** An illustration of the environment for *n* = 3 and a single example rule. **(B)** A simplified depiction of a single trial from the perspective of a single agent *m* who is an asocial or belief-based social learner. The steps shown are: **[1]** population receives context *x*_*t*_ **[2]** agent *m* forms lossy representation 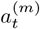 **[3]** agent *m* emits social cue 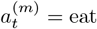 **[4]** if agent *m* is a social learner, they update their beliefs using cue 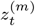 from agent *n*≠ *m* **[5]** agent *m* chooses arm 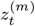 **[6]** iff 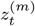, agent *m* updates beliefs with ground truth feedback *y*_*t*_. If agent *m* were instead a *policy-based* social learner, they would simply copy agent *n*’s action directly. **(C)** The distributions over encoder outputs *z*_*t*_ for illustrative values of *κ* and a single example *x*_*t*_

The task proceeds trial-by-trial: at trial *t* ∈ [0, *T* ™ 1] each agent *m* in the population has fitness 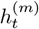, receives *shared* context *x*_*t*_ ∈ *X*_*n*_^1^, independently selects arm *a*^(*m*)^ ∈ {eat, discard} and receives reward 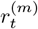 according to

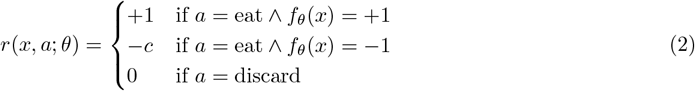

i.e. there is a ‘safe’ arm (that always produces zero reward) and a ‘risky’ arm that produces rewards of +1 or ™*c* for different contexts. The fitness of each agent in the population is then updated as

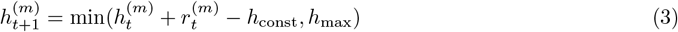

where *h*_max_ is the maximum achievable fitness and *h*_const_ is the baseline rate of fitness loss. If 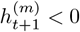, agent *m* ‘dies’ and is removed from the environment. Unless otherwise specified, we will use *c* = 1, *h*_0_ = 5, *h*_max_ = 10, *h*_const_ = 0.01 throughout all experiments; see S1 Text for a sensitivity analysis of these and other parameters.

### Asocial learning as Thompson Sampling

Thompson Sampling (TS) is a principled and widely used approach for balancing exploration and exploitation in bandit problems [32, 33], and has also been used to model human behaviour in tasks similar to ours [34, 35]. A TS agent maintains a posterior belief *b*_*t*_(*θ*) over the parameters of their environment; at each trial, they sample a single point 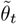 from *b*_*t*_, and then select the arm that maximises expected reward under 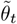. The resulting environment feedback is then used to form updated belief *b*_*t*+1_(*θ*). In our case, each agent in the population maintains an independent belief 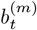(*θ*) about the rule *θ* that controls which species are edible via *f*_*θ*_:

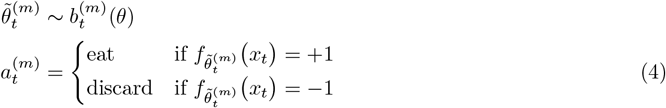

Since our goal is to study the impact of capacity constraints, we must somehow limit agents’ ability to process information. There are multiple ways we could approach this within the TS framework; we do so in terms of the *number of features* (in *x*_*t*_) that agents can perceive during each trial. This choice of implementation is motivated by a focus on *representational* capacity: that is, agents’ ability to represent the task-relevant features of their environment. This is distinct from e.g. functional capacity—regardless of how much information is retained from *x*_*t*_, all agents are equally capable of performing the computations necessary to evaluate rewards, update beliefs, select actions, etc.

Formally, given input *x*_*t*_, we say that each agent forms an individual internal representation 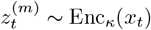 where Enc_*κ*_ preserves on average *κ* bits of information per *x*_*t*_. E.g. when *κ* = 0, agents are unable to process *any* input features; when *κ*≥ *n* agents can process all incoming information; when 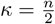 each agent perceives on average half of the features at each trial (see Fig 1C). Note that features are included or excluded independently with uniform probability, so an agent may perceive one feature at trial *t* and a different feature at trial *t* + 1.

Our encoder-based implementation follows work in resource-rational analysis that uses the information bottleneck framework as a principled way to model capacity limits in human and animal cognition [18, 36–39]. In S1 Text, we report results for two related but alternative ways of degrading agents’ access to information; again, these variants constitute different representational bottlenecks rather than an exhaustive survey of possible capacity-related constructs.

Action selection then proceeds on the basis of 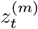 in place of *x*_*t*_, with Eq (4) becoming^2^

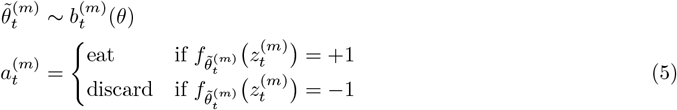

If (and only if) an agent chooses 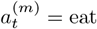, they obtain the ‘ground truth’ category label *y*_*t*_ = *f*_*θ*_(*x*_*t*_) and update their beliefs by suppressing the probability of all rules inconsistent with datapoint 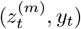:

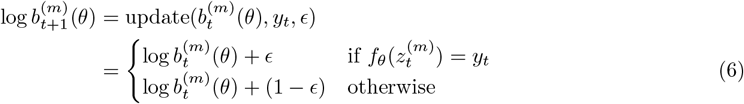

where *ϵ* ∈ [0, 0.5] controls how aggressively beliefs are updated (we will use *ϵ* = 0.05 throughout all experiments)^3^. It is worth emphasising here that the true (uncompressed) feature vector *x*_*t*_ plays no direct role in the agent’s policy: *both* action selection *and* belief revision use only each agent’s lossy representation 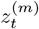, ensuring that *κ* exerts a ‘global’ influence over agents’ ability to learn and make decisions.

As an illustration of how complexity (number of features *n*) and representational capacity (*κ*) interact to determine the difficulty of the asocial learning problem, Fig 2 shows the fitness over time of different capacity-limited agents in the least (*n* = 2) and most (*n* = 6) complex environments. In both environments, we see a consistent positive relationship between capacity and fitness. We also see a significant difference between the two complexity levels: at *n* = 6, even agents with unlimited capacity (*κ* = *n*) are unable on average to reach the fitness ceiling *h*_max_.

**Fig 2.**
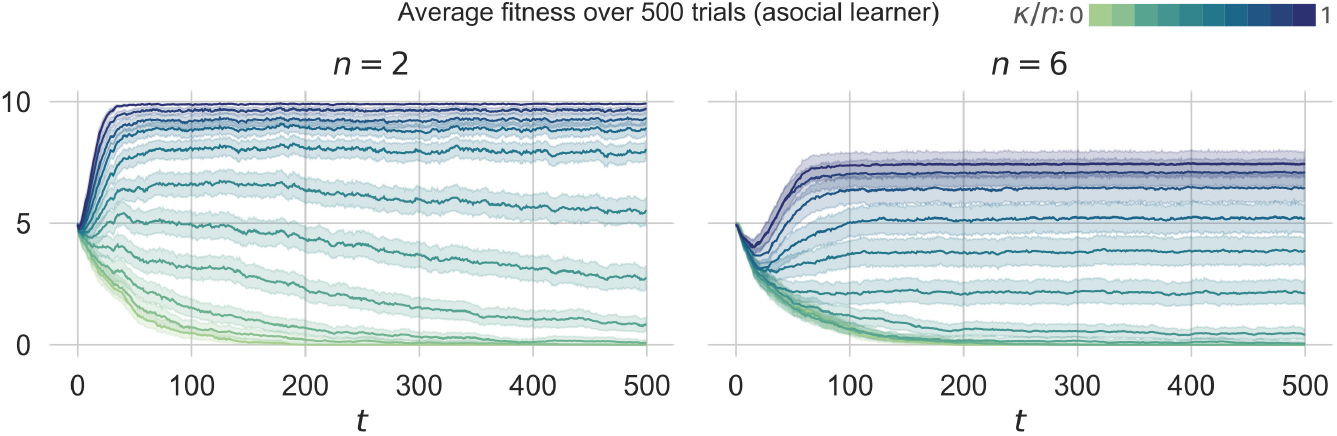
The fitness over 500 trials of a purely asocial learner with different levels of representational capacity (*κ*) in the simplest (left) and most complex (right) task environments. Each line shows the mean over 50 rules sampled uniformly from Θ_*n*_ (with shaded areas giving bootstrapped 95% CIs).

### Adding social information use

Eqs (5)–(6) define our *asocial* learning process. As with capacity, there are a number of different ways we could model social information use, each inviting different cognitive or biological interpretations. We implement two model variants that aim to capture distinct levels of the social learning hierarchy [6, 40, 41]. However, we do not claim that these two variants are exhaustive; generalising our experiments to additional forms of social learning would be a worthwhile direction for future work.

### Policy-based social learning

One of the simplest ways that an agent can learn socially is through direct imitation. In this variant, the social learner simply copies the action choice (eat or discard) from another agent chosen uniformly at random. This represents a maximalist version of social learning, in the sense that it entirely replaces the asocial learning and decision-making processes.

### Belief-based social learning

A more nuanced approach is for social information to influence learners’ beliefs about the world (i.e. *θ*), while action selection still proceeds according to Eq (5). We model this as follows. Immediately after forming representation 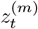 of context vector *x*_*t*_, each agent in the population expresses a *social cue* 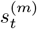 capturing their ‘best guess’ about whether *x*_*t*_ is edible or poisonous:

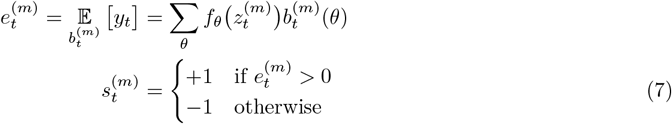

If agent *m* is a belief-based social learner, they then select another agent *n* uniformly at random, and perform a pre-action belief update following Eq (6) as log 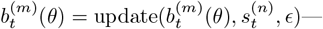that is, social and asocial learning operate under this model via identical ‘mechanics’, with the only difference being that the social belief update uses cue *s* in place of ground truth *y*. Crucially, the ‘social belief update’ takes place before the learner acts, and is independent of which arm is subsequently chosen; it thus captures the idea that social learning provides a complementary source of information that sidesteps the need for direct agent-environment interaction. While we intentionally treat the cue *s* as an abstract signal of a particular agent’s beliefs, it could be interpreted either as something that is *inferred* by the social learner (on the basis of observed actions), or as the content of some explicit communication between agents (such as an utterance or gesture).

Note that for now we are modelling all social learners as using *unbiased transmission*; that is, whether following the policy-based or belief-based approach, they are equally likely to learn from any other member of the population. This approach is also sometimes referred to in the literature as ‘linear imitation’, since the probability of a social learner updating on cue value *s* or copying action *a* scales linearly with the frequency of *s* or *a* in the population. While decades of research have established that humans and other animals are in fact biased social learners, preferring information from some conspecifics over others [5, 6, 22, 42–44], our focus on the *initial* emergence of social information use means that an indiscriminate ‘strategy’ is appropriate. We will relax this assumption in our final two experiments (Sections -).

### Evolutionary dynamics

Our evolutionary simulations are based on the Moran process [45, 46], a simple model of selection in finite populations. In the Moran process, individuals are characterised by a set of fixed types; at each step, a single individual is chosen uniformly at random to die, replaced by a newborn whose parent is selected with probability proportional to their fitness, and who inherits their parent’s type (with some small probability of mutation *p*_mut_)—thus maintaining a constant population size. We use the same “one out, one in” reproductive dynamics, but agents in our simulation die when their fitness falls to zero (with background probability *p*_death_ of dying randomly before then)—so any given timestep may see no deaths, one death, or multiple deaths. We will use *p*_mut_ = 0.01, *p*_death_ = 0.001 throughout our experiments—for a sensitivity analysis of these (and other) parameters, see S1 Text.

In our simulations, each agent’s type corresponds to and determines their learning behaviour: fully-asocial or unbiased-social in Experiment 2 (Section); the same plus conformist-social and copy-fittest-social in Experiments 3-4 (Sections -). In this way we are adopting a form of the phenotypic gambit [47, 48], allowing a single mutation to transform a purely asocial learner into a social information user, or an unbiased social learner into a strategic one. Clearly, this abstracts away a huge amount of complexity; as a simplifying assumption, it is necessary to make practical the kinds of experiments we wish to run.

Since newborn agents inherit no actual *knowledge* from their parents, the overall process can in one sense be viewed as implementing a crude form of meta-learning: in the ‘inner loop’, each agent begins life as a blank slate and must learn the correct mushroom-eating behaviour in order to survive; in the ‘outer loop’, a selection pressure shaped by the combination of complexity and capacity pushes agents towards the most effective learning strategy.

## Results

### Capacity shapes the utility of social information

We first describe the results of a preliminary experiment in which we assess the impact on survival from social information use under different conditions of environment complexity (*n*) and representational capacity (*κ*). We simulate a background population of 100 *asocial-only* learners (i.e. their types are fixed, with mutation disabled) for *T* = 1500 trials; after a warmup period of 500 trials, we ‘drop in’ two new agents—one that uses social information and one that doesn’t—and then record the difference in their subsequent lifespans. This is intended to capture the scenario of a single mutant social learner appearing within an established population of organisms that otherwise learn only via asocial interaction with their environment: how does the capacity of this population to process information determine the fitness advantage enjoyed by the mutant individual? To ensure that our results are robust, we run 500 separate simulations for each (*n, κ*) pair, over rules sampled uniformly from Θ_*n*_.

Earlier, we posed the question of whether the value of social information increases or decreases with a population’s representational capacity. We now offer initial evidence that, to some extent, *both* of these relationships hold true. Fig 3 shows the average excess lifespan for the social learner as a function of *κ*, for each value of *n*. Rather than either a monotonic increase or decrease, we instead observe for both social learning variants an ‘inverse-U’ relationship: at very low *κ* all agent types are ∼ equally bad at surviving; as we increase *κ* the social learner ekes out an advantage, which peaks before decreasing as *κ* approaches *n*. In the case of belief-based social learning (panel A) we also observe an interesting interaction between *n* and *κ*: in low-complexity environments (*n* = 2, 3) there is a region of the curve (0 *< κ* ⪅ *n*2) where social information use is actually *maladaptive*, and the social learner is worse at surviving than their social-information-ignoring counterpart. This effect disappears in more complex environments (*n*≥ 4), where at low capacity the asocial learning problem is so difficult that updating on unreliable cues makes little difference. Interestingly, an equivalent maladaptive region is not observed under policy-based social learning. This difference is explained by the fact that receiving an incorrect cue at trial *t* can “contaminate” the belief-based learner’s posterior, and thus their future behaviour—which can lead them to perform worse than they would have via asocial learning. By contrast, copying an incorrect action at time *t* impacts only the policy-based social learner’s performance on trial *t*, with no downstream consequences. Policy-based social learning may therefore be *ineffective* (i.e. provide zero advantage), but if actions are being copied solely from asocial learners, it cannot be actively maladaptive.

**Fig 3.**
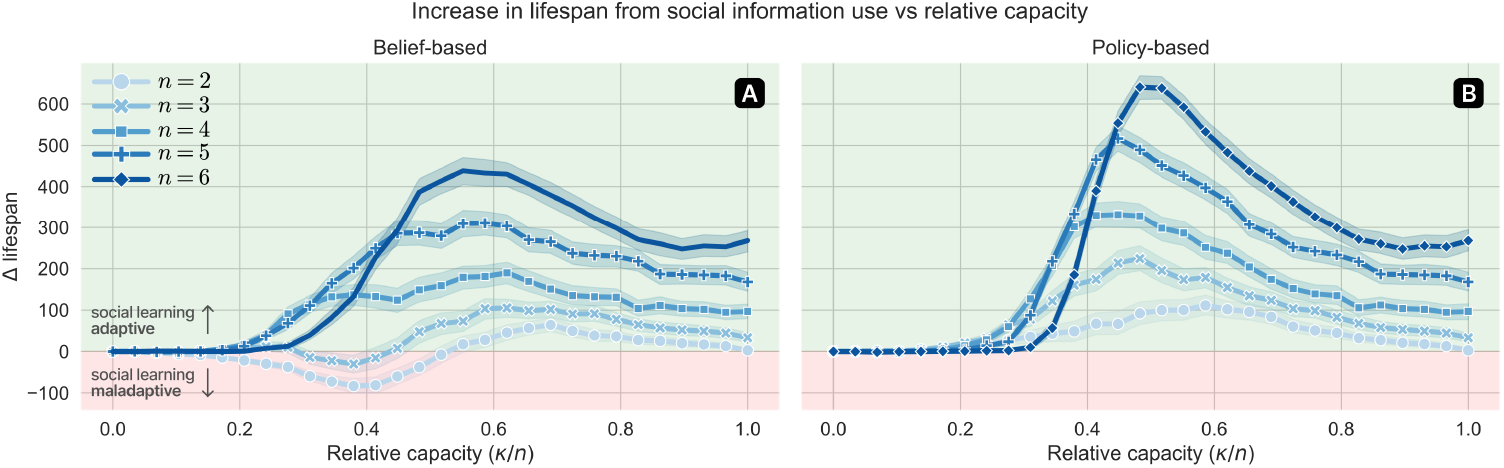
Results of Experiment 1. The difference in lifespan (over 1000 timesteps) for an agent that does/doesn’t engage in policy-based (panel A) or belief-based (panel B) social learning, in a population otherwise composed entirely of asocial-only learners. Each point represents the mean over 500 simulation runs, with rules sampled uniformly from Θ_*n*_ (shaded areas give bootstrapped 95% CIs).

### Capacity shapes the emergence of social learning

To test whether the trend observed in Experiment 1 translates into an effect on the *emergence* of social information use, we now move beyond our analysis of the single-mutant case and allow all agents to occupy either type. Beginning with an initial population of 200 non-social-learners (where social learners can this time appear by mutation), each (*n, κ, θ*) simulation is run for *T* = 10^4^ trials, after which we record the number of social learners in the population. We also run a comparable set of control simulations in which all social information use is disabled so that both agent types have identical learning behaviour. This allows us to estimate the expected type proportions that arise under neutral drift, solely from the effect of capacity on each population’s turnover rate; subtracting these baseline proportions yields an estimate of the actual selection pressure.

As Fig 4 shows, we observe a similar qualitative pattern to that found in Experiment 1: selection for social learners varies roughly as an inverse-U-shaped function of capacity; and is stronger in more complex environments (higher *n*). We also find again that, under certain combinations of *n* and *κ*, belief-based social learning is actually maladaptive. Interestingly—in contrast to Experiment 1—we now also observe a small amount of negative selection in the policy-based case. This is almost certainly the result of a population-level dynamic akin to Rogers’ paradox [22], where a too-high frequency of imitators effectively parasitises the supply of new information. That this effect is weaker for policy-based than belief-based social learning reflects the fact that the latter can be rendered maladaptive by *both* individual-level (belief contamination) and population-level mechanisms. This combination of factors also explains why the maladaptive region for belief-based social learning is larger than in Experiment 1, persisting at higher values of *n*.

**Fig 4.**
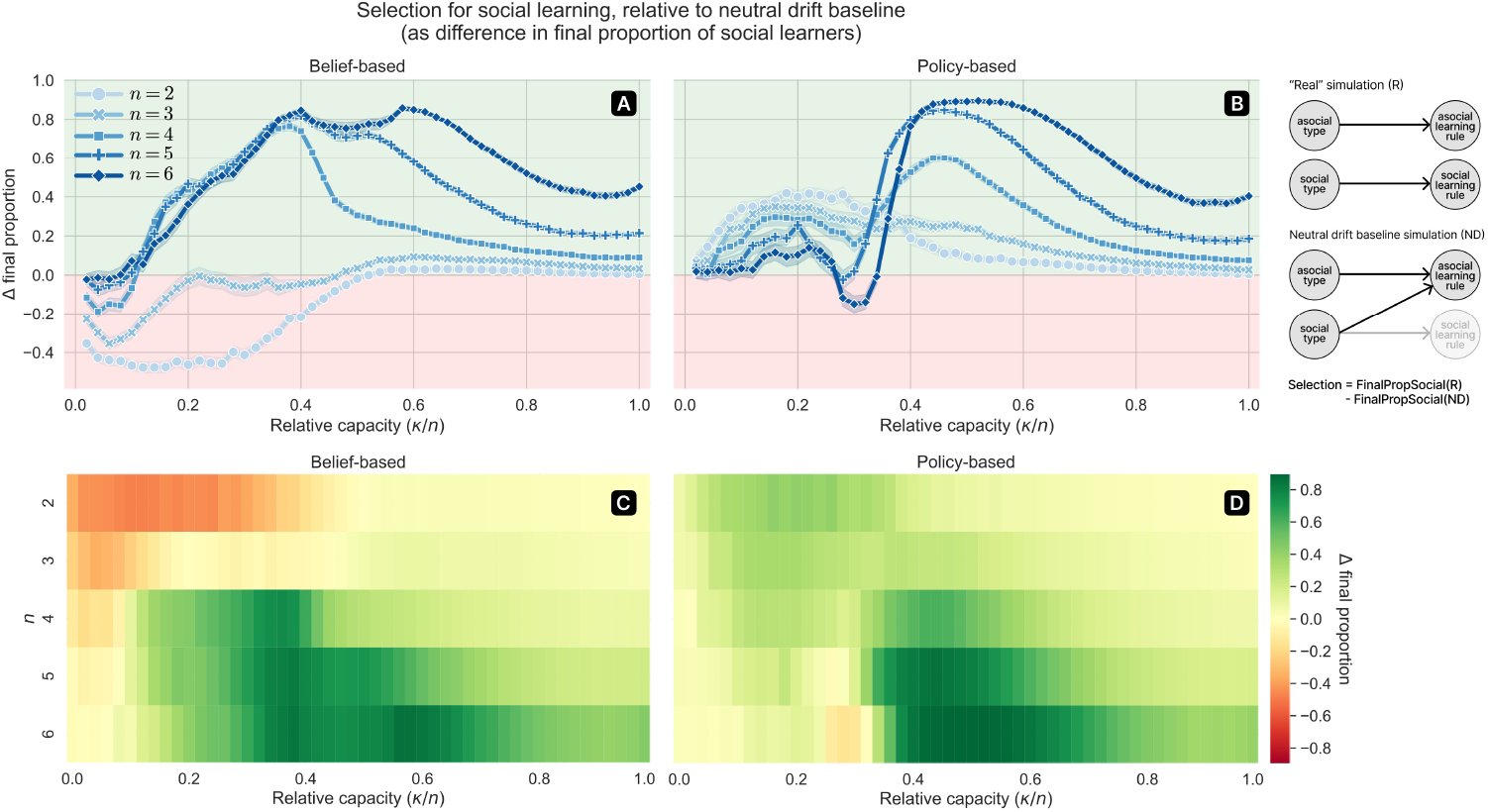
Results of Experiment 2. Empirical selection for social learning under different combinations of complexity (*n*) and capacity (*κ*). Selection is measured as the difference in final population proportion of social learners between simulations where the behavioural effect of the ‘social’ type was enabled (“real” simulation) or disabled (neutral drift baseline). Panels A and C show results for the belief-based social learning variant; panels B and D for the policy-based variant. Each point represents the mean over 500 independent simulation runs, with rules sampled uniformly from Θ_*n*_. All simulations were run for 10^4^ timesteps.

### Capacity shifts the balance between social learning strategies

One limitation of our first and second experiments is their restriction of all social learners to simple unbiased transmission—which arguably describes only a very small minority of the social learning behaviour observed ‘in the wild’ [6, 42, 44, 49]. It would be interesting to try and capture any interactions that capacity limits might have with the *strategies* that human and animal social learners rely on to target their learning towards particular conspecifics.

In our third experiment, we extend the model to accommodate two additional such strategies that are well-studied both empirically and theoretically [5, 50–52]. We implement these strategies as different distributions over potential demonstrators, while otherwise holding fixed the content and mechanics of social learning. That is, all social learners still receive either a cue or action as described earlier; they differ in how they select the source of that cue or action. Formally, each strategy is implemented as a particular distribution over which ‘other’ agent *n*≠ *m* is selected by agent *m* as a source of social information:

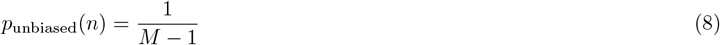

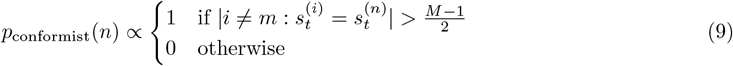

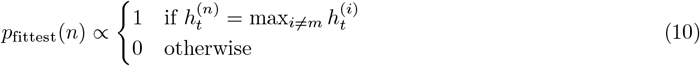

where *M* is the population size and | · | returns the number of elements in a set. Eqs (8)–(10) are given in terms of the belief-based model, but identical formulations are used for the policy-based variant, where all *s* terms are simply replaced with *a* terms. The unbiased learner (Eq (8)) samples uniformly from all its peers, as in Experiments 1-2. The conformist learner (Eq (9)) first identifies which cue or action is expressed by the majority of its peers, and then samples uniformly from agents expressing that value^4^. The copy-fittest learner (Eq (10)) samples uniformly from the currently fittest agent(s) among its peers. These hard-threshold rules should be read as idealised limiting cases of a broader space of strategies in which selection probability varies smoothly with frequency or fitness/success. While our motivation was to avoid introducing additional free parameters, we note that there is some precedent for these formulations [26, 51, 53].

Keeping all else unchanged, we re-ran the evolutionary simulations from Experiment 2, this time using an initial equal split of asocial and unbiased social learners, and focusing on the intermediate-complexity (*n* = 4) environment. This is intended to capture the scenario where ‘strategic’ (i.e. biased) social information use appears by mutation within a population in which unbiased social learning is moderately well established. As before, we report the final proportion of each type in the population following *T* = 10^4^ trials.

Panels B and D of Fig 5 illustrate an interesting result: at lower capacity, social learners using the ‘copy-fittest’ strategy are favoured relative to those using the conformist strategy; at higher capacity the reverse is true. The intuition for this pattern is as follows. When capacity is low, large inter-agent variance in knowledge (and thus fitness) can arise simply from a few individuals getting ‘lucky’—variance which is successfully exploited by the copy-fittest strategy. At higher capacity, where most agents are making predictions above chance, these differences in knowledge are less pronounced, and are outweighed by the insulation that conformity provides against individual noise. To validate this intuition, we estimated the probabilities that strategic belief-based social learners obtain accurate social cues (i.e. 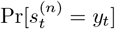; see S1 Text for details on how this is estimated). Similarly, for policy-based social learners, we estimated the probability of each strategy copying the correct action. As suggested by panels B and D, panels C and F confirm that at low capacity these probabilities are highest for copy-fittest, but that there is a crossover point after which it is always highest for conformity. Strikingly, this analysis further reveals that in the low-capacity regime, simple unbiased transmission is *also* likelier to obtain both accurate cues and correct actions than conformity—suggesting that under some conditions, an apparently naïve approach to social learning can be preferable to a more strategic one.

**Fig 5.**
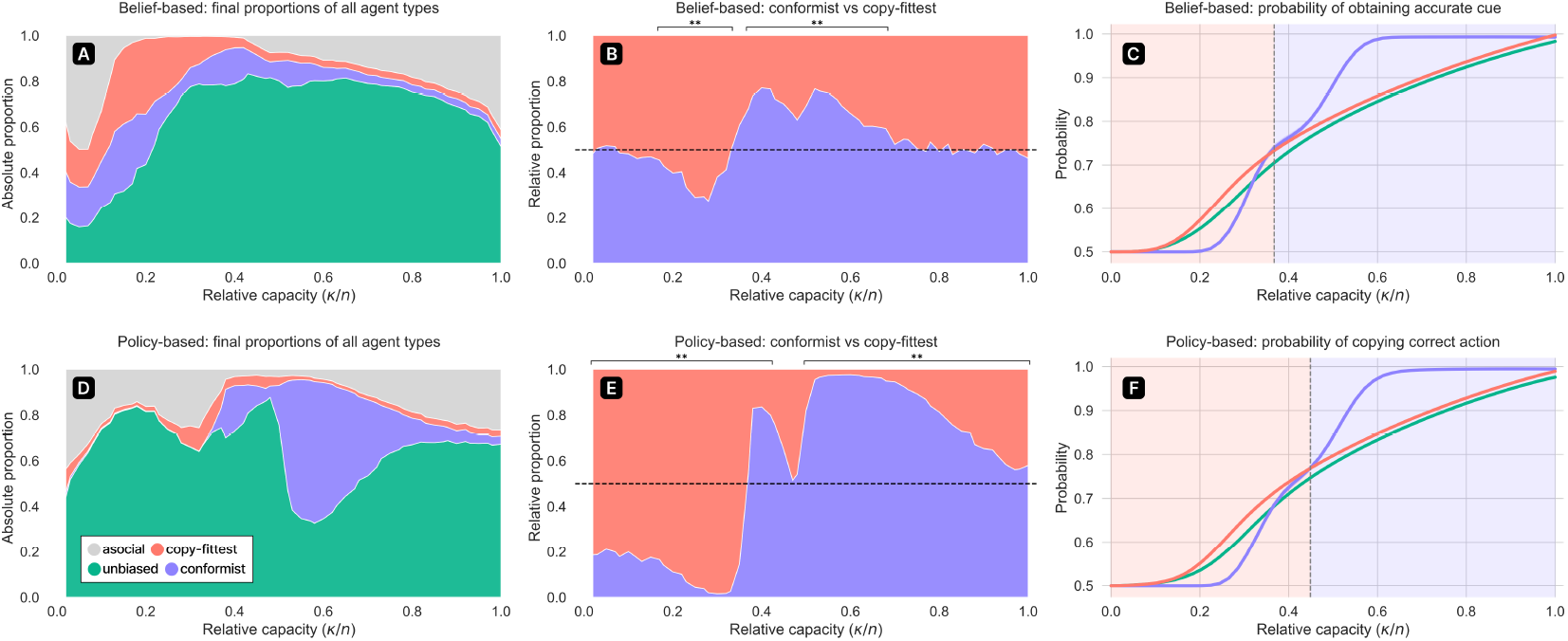
Results of Experiment 3. **(A)** the final proportion of each agent type in the population after 10^4^ timesteps of evolutionary simulation under different values of capacity (*κ*), in the *n* = 4 environment and using the belief-based social learning model. Proportions were aggregated over 500 independent simulation runs, with rules sampled uniformly from Θ_4_. **(B)** from the same data, the relative proportions of conformist and copy-fittest strategies. Brackets indicate *p*-values *<* 0.01, using a Wilcoxon signed-rank test with a null hypothesis of equal proportions. **(C)** the estimated probability that the cues obtained by each social learning strategy are accurate (i.e. *s*_*t*_ = *y*_*t*_), based on a simulation where no social learning actually occurs. Shaded regions denote timesteps at which copy-fittest learners outnumber conformists and vice versa. Panels **D**-**F** are equivalent to **A**-**C** but for the policy-based social learning model.

As in previous experiments, it is worth noting that we again see a difference between the belief-based and policy-based variants; with relative selection for both strategies, and absolute selection for conformism, considerably stronger in the policy-based case. One plausible explanation for this is simply that the difference between conformist and copy-fittest strategies is sharper in the policy-based model, where the impact of social information on behaviour is much more *direct*. The more roundabout nature of the belief-based approach might mean that the benefits of either strategy are softened—especially at low capacity.

### Learning strategies co-evolve with capacity

All of the evolutionary simulations reported thus far have held each population’s representational capacity constant, while allowing only agents’ learning types to evolve. The results of Experiment 3 suggest that, within a population whose capacity evolves upwards over time, and which initially contains no ‘strategic’ social learners, we should expect to see two separate transitions; where copy-fittest social learners first obtain a majority before themselves being overtaken by conformists. For our final experiment, we test this prediction by jointly evolving both capacity and agent types within a single population. To capture the intuition that capacity, as a more ‘fundamental’ attribute of cognition, should evolve more slowly than specific learning strategies, we allow *κ* to mutate only every 20 timesteps:

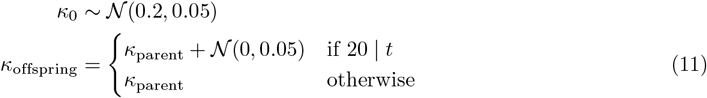

The results of Experiment 4 are shown in Fig 6. As we would expect, mean capacity (panels A,D) increases monotonically over time, while panels B,C,E,F confirm that we do indeed observe the two predicted transitions. Initially, copy-fittest agents are more effective at invading the population. These dynamics are then steadily reversed, with conformists soon gaining a plurality that is retained for the remainder of the simulation time. Interestingly, the frequency of asocial learners crashes to ≈ 0 almost immediately in the belief-based model (panel B), but follows a much steadier decline in the policy-based model (panel E). This is almost certainly a consequence of the same Rogers’-paradox-like dynamics that contributed to negative selection in Experiment 2. The difference arises because belief-based social learners are not pure free-riders; while they use information from others to update their beliefs, they still choose actions and emit their own cues independently. By contrast, policy-based social learners *are* pure free-riders, contributing no independent information back to the common pool. Asocial learners are thus more valuable to the population under policy-based than belief-based social learning; leading us to observe a much stronger frequency-based effect in the former case.

**Fig 6.**
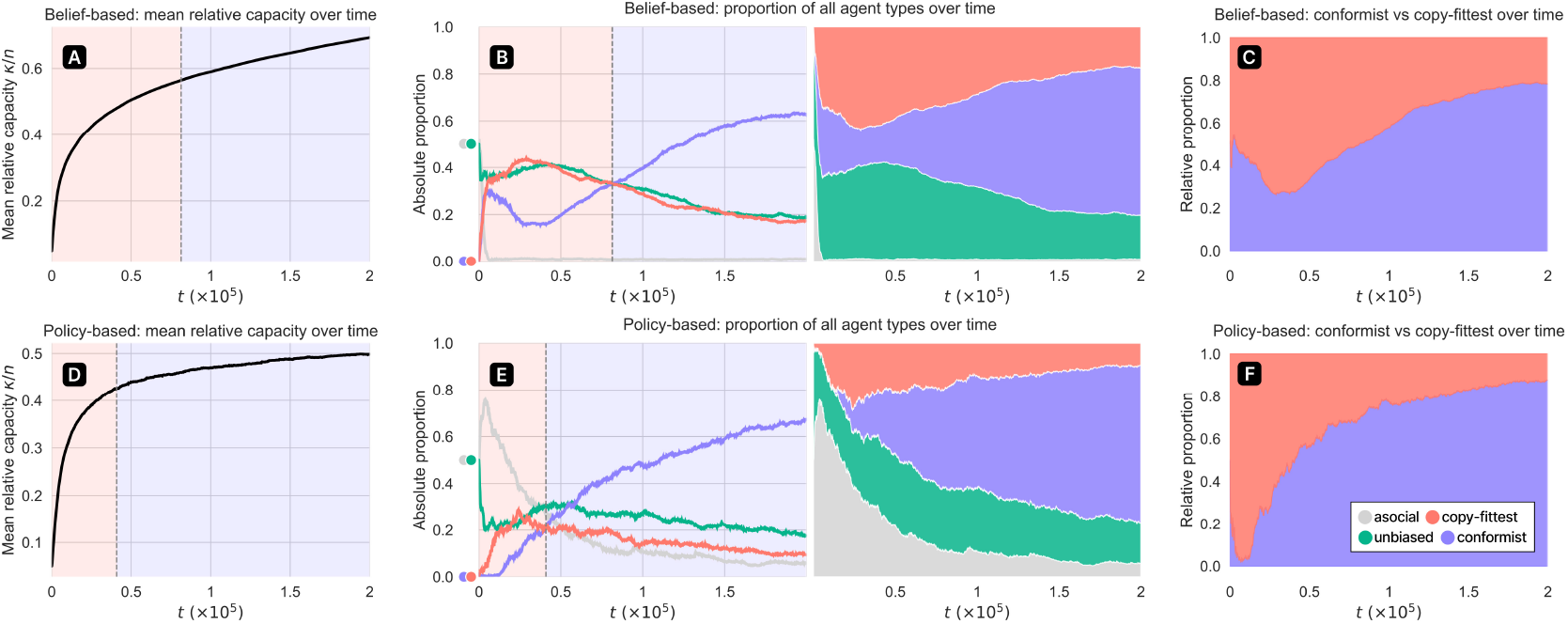
Timeseries data from agents undergoing simultaneous co-evolution of representational capacity and learning type, in the *n* = 4 environment. **(A)** The change in average capacity over time. **(B)** The proportion of each agent type in the population over time. **(C)** The *relative* proportions of conformist and copy-fittest social learners over time. All plots report mean values over 200 independent simulation runs, with rules sampled uniformly from Θ_4_. Shaded regions of panels A-B denote timesteps at which copy-fittest learners outnumber conformists and vice versa.

## Discussion

Many animals, either in infancy or as a response to migration or environmental change, must at some point learn about an unfamiliar world while avoiding the various dangers that it poses. Faced with this dilemma, social learning offers a valuable way to acquire adaptive behaviours while mitigating the cost and risks of trial-and-error exploration. In understanding the factors that strengthen or weaken this effect, previous work has studied in depth the role of environment properties such as temporal variability [5, 22–25] and task difficulty [26, 27]. Where consideration has been given instead to the impact of *agent* -centric properties, it has mostly been in terms of different social learning strategies, balances between social and asocial learning, or network structures [6, 28–30]. Less attention has been given to the role of more ‘fundamental’ or domain-general properties of cognition—which, by virtue of being slower to evolve, ought to exert influence over the emergence of narrower adaptations such as whether or how learners integrate social information.

In this paper, as a step towards a broader understanding of how agent and environment properties interact to determine which learning strategies are most successful, we have explored how the evolutionary benefits of social learning depend on agents’ ability to represent task-relevant information. Our focus on capacity is motivated by the resemblance of the cost-avoidance argument to ideas in resource-rational analysis [15].

While this resemblance has led some to characterise social information use as a resource-rational solution to the problem of learning in complex and dangerous environments [19–21], the relationship between information-processing capacity and the utility of social learning had not previously been tested.

Such an exploration may initially seem redundant: decreasing an agent’s ability to process or represent information is surely equivalent to increasing the difficulty of the asocial learning problem, and work such as [26, 27] already tells us what to expect as a result. But this is only true if we assume that the learning agent’s *sources* of social information are not similarly constrained. Since any realistic treatment must reject this assumption, the picture becomes less straightforward: on one hand, decreasing capacity makes the asocial learning problem harder, which should favour social learning; on the other hand, it reduces the quality of social information, which should favour asocial learning. Through a series of simulation experiments, we have shown that the combined result of these two effects is that the adaptiveness of social learning follows an ‘inverse-U’ relationship with capacity, decreasing towards both extremes of the spectrum.

How should this result be interpreted? Whether social learning operates as an auxiliary information source complementary to the primary channel of asocial feedback (as in our belief-based model) or as a surrogate policy that replaces individual exploration (as in our policy-based model), its value will depend on the balance of two factors: the amount of reliable information conveyed, and the extent of the learner’s marginal need for such information. The first of these factors increases with population capacity: higher *κ* means that demonstrators express more accurate cues and take more adaptive actions (and in the belief-based model, cues can furthermore precipitate larger belief updates). Meanwhile, the learner’s informational need decreases with capacity—as *κ* approaches *n*, the asocial learning problem becomes more straightforward, reducing the space available for improvements in fitness from access to extra information.

The combined effect is that social learning is most useful when capacity is *high* enough that social information is moderately informative but *low* enough that the learning problem is still difficult (see Table 1).

**Table 1.** A summary of the effect of representational capacity on the two factors that determine the value of social information use.

| $\kappa/n$ | social information quality | learner need |
| --- | --- | --- |
| low | low | high |
| moderate | moderate | moderate |
| high | high | low |

While our main qualitative findings held across both belief-based and policy-based social learning, each of our experiments surfaced differences between the two variants—highlighting some interesting consequences of how social information is integrated into downstream behaviour. Perhaps the most interesting of these is that belief-based social learning was more susceptible to “individual-level” maladaptivity (via contamination of future choices) whereas policy-based social learning was more susceptible to frequency-dependent effects in the vein of Rogers’ paradox. Future work may explore connections to recent findings from Roberts-Gaal et al. [54], who show that ecological stability influences the format of cultural learning by shaping a tradeoff between efficient but inflexible procedural information and more flexible but costly goal- or causal-structure information. This difference in “failure modes” between our two social learning variants highlights a related tradeoff; namely that more flexible, less localised integration of social information can also increase the downstream consequences of exposure to unreliable demonstrators.

Also of relevance to our results is the work of Fogarty et al. [55], who studied the impact of task difficulty on the evolution of explicit costly teaching, as opposed to ‘inadvertent’ social learning. Notably, the authors also found an “n-shaped function” where teaching was not favoured for traits that were either too easy to learn (where learners can readily acquire them through other means) or too difficult (where teachers are unlikely to possess the relevant information). One way to view our findings is that they demonstrate how Fogarty et al.’s results for explicit teaching can hold also for inadvertent social learning, given a construction of task difficulty that arises from constraints on individuals’ representational capacity.

### Limitations and directions for future work

While our current models are focused on the relative adaptiveness of social vs asocial learning (and between different social learning biases), it is important to recognise that resource constraints also influence the strategies that agents deploy during asocial learning itself. For instance, constraints on executive function have long been known to shift human learners away from computationally expensive model-based planning and toward simpler, model-free heuristics [56]. More recent work has shown, similarly, that human learners will limit (costly) uncertainty-directed exploration in favour of (cheaper) random exploration under increases in working memory load [57], time pressure [58] or search space complexity [59]. This line of work not only offers a potential alternative formulation of capacity in terms of ability to explore, but also suggests an additional route by which generic capacity limits might favour social learning (as an alternative to costly directed-exploration strategies). Explicitly connecting our findings to richer and more structured forms of asocial exploration thus represents another promising direction for future research.

A more significant limitation of the current work is that our models do not consider any cognitive costs specific to social learning itself—capacity limits manifest only in agents’ ability to represent their environment *in general*, and have no impact on how successfully they obtain the actions or cues of others.

Doing things this way allows us to isolate how representational capacity changes the supply of and demand for social information. It is, however, an oversimplification, and future work should investigate the effect of introducing social-learning-specific costs. We suspect that doing so would not fundamentally change the inverse-U dynamics found in our experiments, but would likely narrow the adaptive region by rendering social learning less effective at lower capacity values. Such considerations would be particularly salient under an interpretation of our belief-based social learning variant in which belief cues are not directly observable but are instead inferred from actions or similar evidence by a theory-of-mind-like process [40, 60]. The same could be said for a more explicitly pedagogical setting where learner and demonstrator alike must make inferences about one another’s knowledge and intentions [61, 62].

Our comparison between conformist and success-biased social learning also leaves aside any costs of implementing these strategies. In our simulations, learners can always identify the majority or currently fittest individual without error. Success-biased learning in the wild, however, requires estimating which individuals are competent on the basis of noisy, delayed, and/or domain-specific cues. Similarly, conformist learning involves sampling enough demonstrators to estimate the majority, and so may be costly when social access is limited or behavioural frequencies are difficult to observe. Taking such costs into account may shift the two strategies’ relative advantage.

Finally, our foraging environment encapsulate a specific class of learning problem—inferring the discrete (deterministic) rule that explains a sequence of sampled experiences. An alternative approach would have agents generalise from prior encounters based on some measure of perceived similarity between different mushrooms (feature vectors). Rule-based and similarity-based generalisation involve different tradeoffs: rules allow for rapid transfer to novel stimuli, while similarity-based representations can capture a wider space of functional relationships [63]. Future work might test how adopting the latter in place of our symbolic rule inference model affects selection for social learning. For example, it is possible that agents’ asocial learning performance would degrade more gracefully under limited feature access, potentially narrowing the region over which social learning is adaptive, or allowing the maladaptive effect to persist at higher environment complexities.

## Conclusion

In summary, when it comes to the usefulness of cultural learning, not all agents are created equal. Information-poor populations are unable to support reliable social learning, while information-rich populations don’t really need it; only when agents lie within a goldilocks zone of relative capacity does social learning reliably take root. A possible biological implication of our results, which future work should seek to test, is that species most strongly reliant on social learning should be found disproportionately at an intermediate level of representational capacity relative to their ecological demands. Crucially, we suggest that species below this threshold may fail to evolve social learning not because they lack either the need for cheaper, less risky alternatives to trial-and-error exploration, or the ‘machinery’ for integrating social information—but because their conspecifics do not generate cues reliable enough to be worth making use of. This is analogous to observations that cumulative culture emerges only once distinct thresholds of individual problem-solving are breached [64, 65].

But what about humans? We are extremely cognitively sophisticated, and yet each of us typically learns a huge fraction of our skills and knowledge from parents, teachers, mentors and peers: does this not contradict our central message? Not necessarily. While it’s true that humans enjoy highly developed brains, we also continuously modify our environment to create conditions of increasing ecological and cultural complexity. Once social information use emerges, it can play into a feedback loop of cultural niche construction [66, 67], where the cultural spread of information produces more complex environments, which in turn keep humans within the adaptive regime of *relative* capacity and reinforce the demand for more social learning.

Building upon our main finding (the “whether”), we also demonstrated through an additional set of simulations evidence for an effect on the relative adaptiveness of different social learning *strategies* (the “how”). Specifically, we found that learners who selectively use information from only the fittest agents performed well in lower-capacity populations, whereas conformist learners that simply followed the majority opinion were most successful in higher-capacity populations. We offer the following intuition for this result. In a capacity-poor population, a few individuals experiencing a ‘lucky run’ of trials can produce significant variance in knowledge and therefore fitness—in other words, the expected difference in social information quality between the fittest and median-fitness agents is relatively high. In a capacity-rich population, where the majority of agents are above chance, these differences are dampened, and are outweighed by the protection against individual noise that conformity provides.

An analysis of each strategy’s probability of obtaining correct social cues or optimal actions revealed something else surprising: though always worse than copy-fittest, at sufficiently low capacity the naïve approach of selecting a source at random (unbiased transmission/linear imitation) yields better odds than conformist social learning. This stands in contrast with predictions from prior work that conformist social learning should always outperform unbiased transmission [26], and future work should seek to explore this result in greater detail.

## Supporting information

**S1 Text. Supplementary simulation results for alternative capacity variants, robustness analyses, and additional methodological details**.

## Acknowledgments

This section is intended only for general acknowledgements and thanks. Any information related to funding, data availability, author contributions, etc. should be entered directly into their dedicated fields in the PLOS Editorial Manager submission system, which will then be incorporated into the appropriate section in your article during the production process.

## Footnotes

1 Rather than sampling *x*_*t*_ uniformly from *X*_*n*_ we apply a class-balanced correction, so that *p*(*x*_*t*_ edible |*θ*) = *p*(*x*_*t*_ poisonous| *θ*) = 0.5. This ensures that the survival challenge posed by a given environment is purely determined by the difficulty of the *learning* problem (a function of *n* and capacity), not by how hazardous it would be for a randomly acting agent.

2 We reuse the *f*_*θ*_(*·*) notation here for convenience, though technically the function that agents compute is slightly different to Eq (1), as it ignores elements of *θ* that line up with zeroed-out entries in 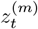.

3 Note that the use of representation 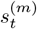 in Eq (6) means that a given rule *θ* might make an incorrect prediction for *x*_*t*_ but still not be downweighted, if the features responsible for the mismatch are not present in 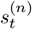

4 Note that since there are only two possible values for both cues and actions, and we always use an even-numbered *M* (and therefore odd *M ™* 1), the conformist strategy doesn’t need to handle the possibility of there being no majority.

