## Supplementary material for "Representational capacity shapes both the “whether” and “how” of social learning": S1 Text

### Supplementary Materials for
### Cognitive capacity shapes both the "whether" and "how" of social learning

#### Contents

**1 Alternative formulations of capacity limits** **S2**
1.1 Decaying beliefs . . . S2
1.2 Noise in reward and cue integration . . . S3

**2 Sensitivity analysis** **S3**
2.1 Experiment 2 . . . S5
2.2 Experiment 3 . . . S6

**3 Other experimental / modelling details** **S6**
3.1 Strategy reliability analysis . . . S6

### 1 Alternative formulations of capacity limits

To ensure that our results are not simply an artefact of the particular way we model capacity constraints (i.e. via partial observation of context features), we implement two alternative model variants and use them to reproduce our main simulation results (Experiments 2-3) for belief-based social learning.

#### 1.1 Decaying beliefs

One way of modelling capacity is to do so in terms of memory. With this approach, the idea is simply that agents with higher capacity should be better able to retain information from previous trials. We implement this through a stochastic process that, at the beginning of each trial, decays agents' current beliefs by some amount towards the uniform distribution.

$$\alpha_i \sim \text{Uniform}(\lambda, 1),\; i = 1 \ldots |\Theta_n|$$
$$\log b_t^{(m)}(\theta_i) = \alpha_i \log b_t^{(m)}(\theta) \tag{S1}$$

where $\lambda \in [0, 1]$ is our measure of capacity. The effect of this is that an agent's current beliefs at any trial are dominated mostly by their recent experiences, rather than shaped equally by their whole history.

Fig AA shows the results from running Experiment 2 with this capacity variant. Importantly, we reproduce the inverse-U shape observed in our original results—as well as the finding that social information use is maladaptive under some parameter combinations (although here for $n > 2$ rather than $n < 4$ as previously). Interestingly, we do not reproduce the finding that social learners have greater advantage in more complex environments (i.e. increasing $n$); on the contrary, the highest area under the curve here is seen for $n = 2$. It is not immediately clear why this is the case; we leave analysis of this for future work.

Fig AB shows the results from running Experiment 3 with this capacity variant. We reproduce the finding that the copy-fittest strategy is favoured at lower capacity and the conformist strategy at higher capacity—although we note that the early invasion of copy-fittest agents is significantly stronger (and the later invasion of conformist agents somewhat weaker) than observed in our original results. One possible explanation for this is that the emphasis on recent experience induced

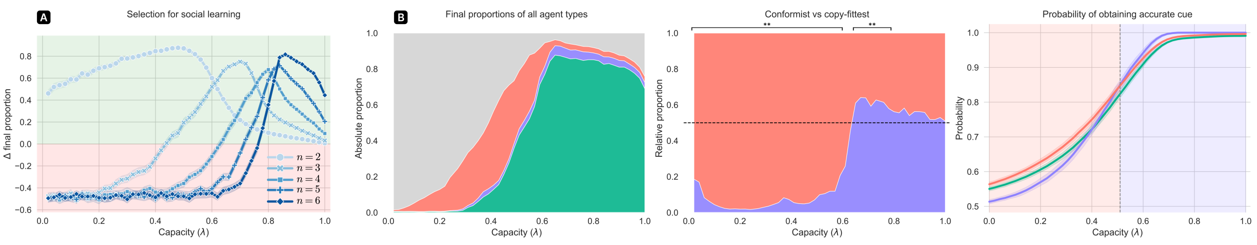


**Fig A in S1 Text**: **(A)** Replication of Experiment 2 for the belief decay variant of our model (Fig 4 in the main text). **(B)** Replication of Experiment 3 for the belief-decay variant of our model (Fig 5 in the main text). Aside from the different implementation of capacity, all other simulation and analysis details remain unchanged from the original version.

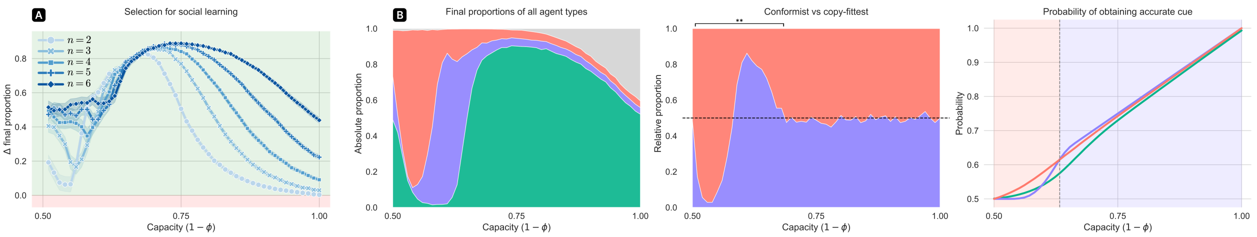


**Fig B in S1 Text**: **(A)** Replication of Experiment 2 for the reward-integration-noise variant of our model (Fig 4 in the main text). **(B)** Replication of Experiment 3 for the belief decay variant of our model (Fig 5 in the main text). Aside from the different implementation of capacity, all other simulation and analysis details remain unchanged from the original version.

by decaying beliefs allows for wider variance to emerge between individuals, which is exploited by the copy-fittest strategy.

#### 1.2 Noise in reward and cue integration

Another way to model capacity, more conceptually related to our original approach, is to introduce information loss not in agents' perception of input features, but in their integration of social and asocial feedback. To implement this, we flip the reward and cue signals used to update agents' beliefs, with some probability $\phi \in [0, 0.5]$. Let $r_t'^{(m)}$ and $s_t'^{(n)}$ be the reward and social cue actually used to update the beliefs of agent $m$ at time $t$. Then,

$$\Pr[r_t'^{(m)} = r] = \begin{cases} 1-\phi, & \text{if } r = r_t^{(m)} \\ \phi, & \text{if } r = -r_t^{(m)} \\ 0, & \text{otherwise} \end{cases} \tag{S2}$$

$$\Pr[s_t'^{(n)} = s] = \begin{cases} 1-\phi, & \text{if } s = s_t^{(n)} \\ \phi, & \text{if } s = -s_t^{(n)} \\ 0, & \text{otherwise} \end{cases} \tag{S3}$$

and we measure capacity as $1 - \phi$.

Fig BA shows the results from running Experiment 2 with this capacity variant. We reproduce once again the inverse-U shape from our original and belief decay variants. In contrast to the belief decay variant, here we do find that the emergence of social learning increases with environment complexity ($n$), as in our original results.

Fig BB shows the results from running Experiment 3 with this capacity variant. Again, we reproduce the 'crossover' from copy-fittest to conformist social learning as capacity increases (with the relative advantage being more evenly matched than seen with our other model variants).

### 2 Sensitivity analysis

As an additional check on the robustness of our main findings, we report the results of a sensitivity analysis over the key parameters of our evolutionary simulations, for our original capacity variant

| Parameter | Description | Default value | Minimum | Maximum |
|---|---|---|---|---|
| $c$ | Fitness cost from consuming a poisonous mushroom | 1.0 | 0.5 | 16.0 |
| $h_{\text{max}}$ | Maximum agent fitness | 10 | 5 | 160 |
| $h_{\text{const}}$ | Background fitness loss per timestep (regardless of action) | 0.1 | 0.05 | 1.6 |
| $p_{\text{death}}$ | Per-agent probability of dying at timestep where $h_t^{(m)} > 0$ | 0.001 | 0.0005 | 0.016 |
| $p_{\text{mut}}$ | Probability that newborn agent has their phenotype assigned at random (rather than inheriting from their parent) | 0.01 | 0.005 | 0.16 |

**Table A in S1 Text**: Evolutionary parameters that were varied for the sensitivity analysis.

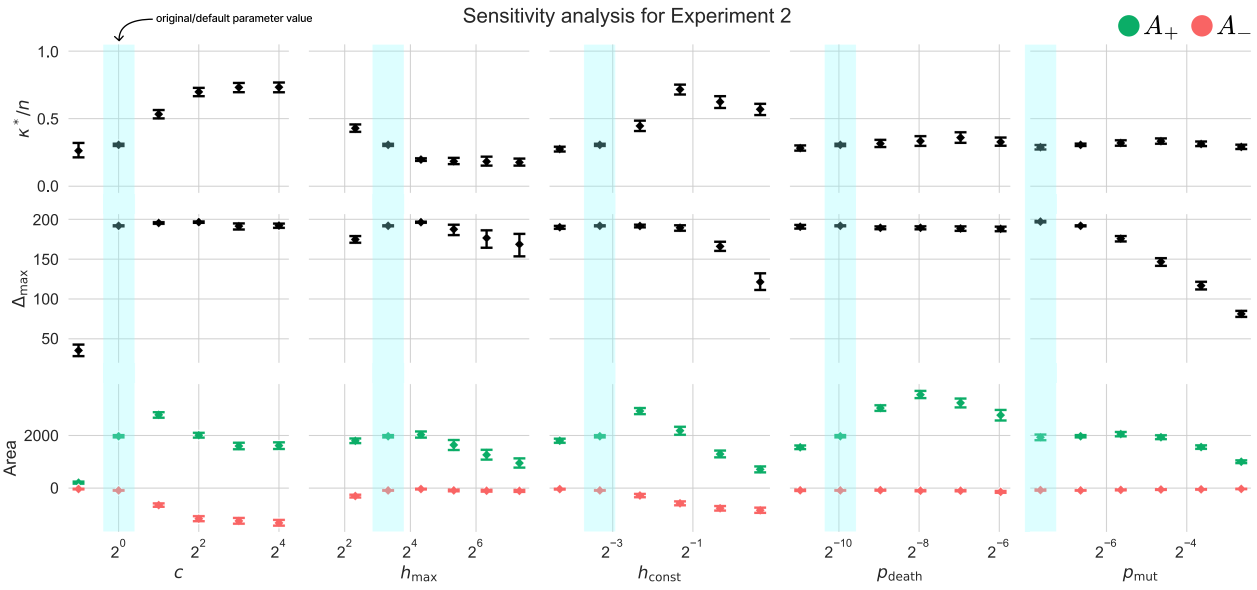


**Fig C in S1 Text**: Results of our sensitivity analysis for Experiment 2. **Top:** capacity value for which the highest number of social learners evolved (relative to control simulation). **Middle:** the final number of excess (i.e. relative to control) social learners in the population for capacity $\kappa^*$. **Bottom:** the total positive (green) and negative (red) difference in social learner counts relative to control simulations, summed over all capacity values (indicating the strength of positive and negative selection respectively). All plots show the mean and bootstrapped 95% confidence intervals over 60 simulation runs, with rules sampled uniformly from $\Theta_4$. Light-blue shading indicates the parameter values used in our original experiments.

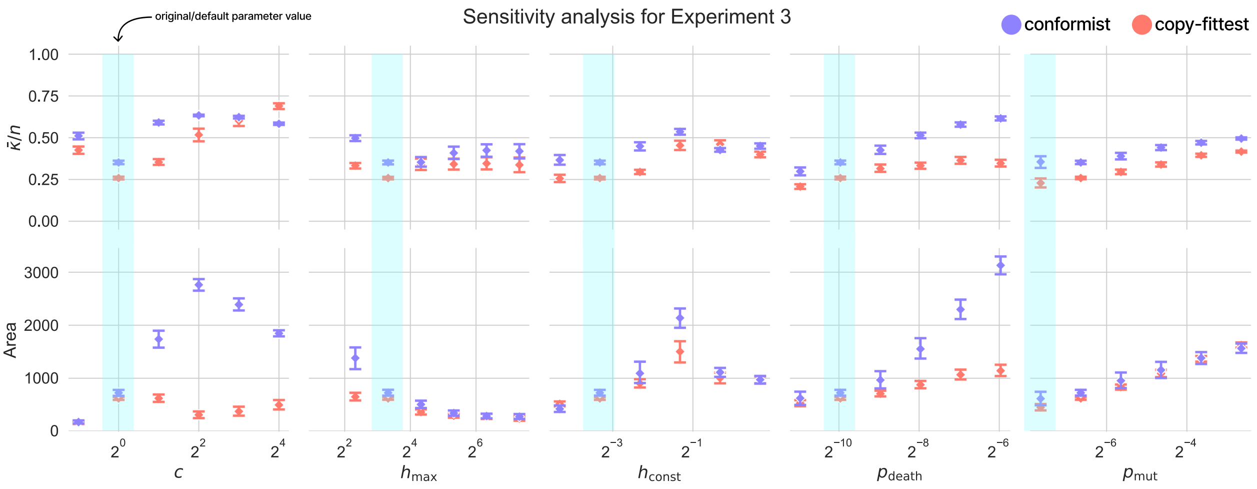


**Fig D in S1 Text**: Results of our sensitivity analysis for Experiment 3. **Top:** capacity centroid for each of the two strategic social learner types. **Bottom:** the total number of agents of each type in the final population, summed over all capacity values. All plots show the mean and bootstrapped 95% confidence intervals over 60 simulation runs, with rules sampled uniformly from $\Theta_4$. Light-blue shading indicates the parameter values used in our original experiments.

(partial observation of features), in the intermediate-complexity ($n = 4$) environment and using belief-based social learning. For each parameter listed in Table A, we reran our main simulations (for Experiments 2-3) over a range of values increasing by factors of 2 from half of the original ('default') value. During each simulation run, all other parameters were fixed to their default values. For compactness, we report a selection of summary metrics intended to capture the most relevant information for each Experiment.

#### 2.1 Experiment 2

For a given setting of evolutionary parameter values $\mathbf{p}$ and capacity $\kappa$, let $\Delta(\mathbf{p}, \kappa)$ be the 'excess' number of social learners in the population after evolution under $(\mathbf{p}, \kappa)$ relative to a control simulation with no social information use (corresponding to the red lines in Figs AA and BA). We then define

$$\Delta_{\max}(\mathbf{p}) = \max_{\kappa} \Delta(\mathbf{p}, \kappa) \tag{S4}$$

$$\kappa^*(\mathbf{p}) = \arg\max_{\kappa} \Delta(\mathbf{p}, \kappa) \tag{S5}$$

$$A_+(\mathbf{p}) = \sum_{\kappa} \max\big(\Delta(\mathbf{p}, \kappa), 0\big) \tag{S6}$$

$$A_-(\mathbf{p}) = \sum_{\kappa} \max\big(-\Delta(\mathbf{p}, \kappa), 0\big) \tag{S7}$$

i.e. $A_+$ and $A_-$ give the total area of the adaptive and maladaptive regions of the curve, respectively. Fig C shows the values of these four metrics over the parameter ranges given in Table A. For assessing the robustness of our original findings, $\kappa^*/n$ (top row) is the most important measure. Values in the middle range would indicate an inverse-U shape for the prevalence of social information use vs capacity—whereas values very close to 0 or 1 would imply that the number of social learners instead simply decreases or increases with capacity. Examining Fig C, we see that all mean $\kappa^*/n$ values are between 0.2 and 0.75. This tells us that the inverse-U trend is preserved across our tested parameter ranges, and thus the conclusion drawn from Experiment 2 is sound.

#### 2.2 Experiment 3

Let $\text{Count}(\pi, \mathbf{p}, \kappa)$ be the number of agents of type $\pi$ in the population after evolution under $(\mathbf{p}, \kappa)$. We then define

$$A(\pi, \mathbf{p}) = \sum_{\kappa} \text{Count}(\pi, \mathbf{p}, \kappa) \tag{S8}$$

$$\hat{\kappa}(\pi, \mathbf{p}) = \frac{1}{A(\pi, \mathbf{p})} \sum_{\kappa} \kappa \cdot \text{Count}(\pi, \mathbf{p}, \kappa) \tag{S9}$$

i.e. a high (low) value of $\hat{\kappa}(\pi, \mathbf{p})$ means that strategy $\pi$ was favoured more under high (low) values of population capacity, for evolutionary parameter values $\mathbf{p}$. Fig D shows the values of these metrics over the parameter ranges given in Table A. Importantly, it shows that the central result of Experiment 3 (copy-fittest social learning being favoured at lower capacity and conformist at higher) holds for the vast majority of tested parameters—with the only two reversals being found at the fairly extreme parameter values of $c = 16$ and $h_{\text{const}} = 0.8$.

### 3 Other experimental / modelling details

#### 3.1 Strategy reliability analysis

Here we explain how the probability estimates plotted in Figs AB and BB (and main text Fig 5C) are obtained. We will first go through the procedure for our original capacity variant (lossy feature encoding), and then briefly describe how things differ for the two additional variants explored above. We will lay out each procedure for the belief-based social learning model only; the analysis for the policy-based social learning model is very similar.

##### Encoder variant

First we must compute the probability, at trial $t$, that agent $i$ emits a social cue correctly tracking the ground truth label $y_t$—that is, $\Pr[s_t^{(i)} = y_t \mid x_t]$. We can write this as

$$\Pr[s_t^{(i)} = y_t \mid x_t] = \begin{cases} \gamma_t^{(i)}(x_t) & \text{if } f_\theta(x_t) = +1 \\ 1 - \gamma_t^{(i)}(x_t) & \text{if } f_\theta(x_t) = -1 \end{cases} \tag{S10}$$

where $\gamma_t^{(i)}(x_t) := \Pr[s_t^{(i)} = +1 \mid x_t]$—that is, the probability that agent $i$ emits a 'positive' social cue (signalling edibility) for context $x_t$. Following Eq (7) in the main text, let

$$e_t^{(i)}(z_t) = \mathop{\mathbb{E}}_{b_t^{(i)}} \left[y_t\right] = \sum_{\theta} f_\theta\big(z_t^{(i)}\big) b_t^{(i)}(\theta) \tag{S11}$$

We then have

$$\gamma_t^{(i)}(x_t) = \frac{1}{|Z|}\sum_{z\in Z}\Pr[z_t^{(i)} = z \mid x_t]\mathbb{1}[e_t^{(i)}(z_t) > 0.5] \tag{S12}$$

where $\mathbb{1}$ is the indicator function, $Z$ is the set of all possible encodings, and $\Pr[z_t^{(i)} = z \mid x_t]$ is determined by the feature alignment between $z$ and $x_t$ and the encoder capacity $\kappa$.

After this, the probabilities of obtaining a correct cue for the *unbiased* and *copy-fittest* strategies are simply given by

$$\Pr[s_t^{(n)} = y_t|x_t, \text{unbiased}] = \frac{1}{M-1}\sum_{i\neq m}\Pr[s_t^{(i)} = y_t \mid x_t] \tag{S13}$$

$$\Pr[s_t^{(n)} = y_t|x_t, \text{copy-fittest}] = \frac{1}{N_{\text{fittest}}}\sum_{i\neq m}\mathbb{1}[h_t^{(i)} = \max_{j\neq m} h_t^{(j)}]\Pr[s_t^{(i)} = y_t \mid x_t] \tag{S14}$$

Where $N_{\text{fittest}}$ is the number of agents $i \neq m$ such that $h_t^{(i)} = \max_{j\neq m} h_t^{(j)}$. For the conformist strategy things are only slightly less straightforward: we need the probability $\Pr[|i \neq m : s_t^{(i)} = y_t| > \frac{M-1}{2} \mid x_t]$. Using for convenience the shorthand $C_t^{(i)} = \mathbb{1}[s_t^{(i)} = y_t]$ we define the discrete random variable $S_t = \sum_{i\neq m} C_t^{(i)}$. We then want

$$\Pr[s_t^{(n)} = y_t|x_t, \text{conformist}] = \Pr[S_t > \frac{M+1}{2}] \tag{S15}$$

$$= \sum_{k=\frac{M}{2}}^{M-1}\Pr[S = k] \tag{S16}$$

Since $S$ follows the Poisson binomial distribution, we can compute this exactly by evaluating $\Pr[S = k]$ for each possible $k$. To obtain the plotted values, we evaluate Eqs (S10)–(S13) averaged over the final 100 timesteps of each simulation run.

##### Belief decay variant

The only difference for the belief decay variant is that, since there are no intermediate representations $z$, Eq (S12) simplifies to

$$\gamma_t^{(i)}(x_t) = \mathbb{1}[e_t^{(i)}(z_t) > 0.5] \tag{S17}$$

##### Reward and cue flip variant

For our final variant, where capacity is taken as the inverse probability of reward and cue signals being flipped, we have the following. Let $p_{\text{unbiased}}(x_t), p_{\text{fittest}}(x_t), p_{\text{conformist}}(x_t)$ be the quantities in Eqs (S13), (S14), and (S15), respectively (with $\gamma_t^{(i)}$ computed as in Eq (S12)). We then simply write

$$\Pr[s_t^{(n)} = y_t|x_t, \text{unbiased}] = (1-\phi)p_{\text{unbiased}}(x_t) + \phi(1 - p_{\text{unbiased}}(x_t)) \tag{S18}$$

$$\Pr[s_t^{(n)} = y_t|x_t, \text{copy-fittest}] = (1-\phi)p_{\text{fittest}}(x_t) + \phi(1 - p_{\text{fittest}}(x_t)) \tag{S19}$$

$$\Pr[s_t^{(n)} = y_t|x_t, \text{conformist}] = (1-\phi)p_{\text{conformist}}(x_t) + \phi(1 - p_{\text{conformist}}(x_t)) \tag{S20}$$

where $\phi$ denotes the probability that cues selected by the social learner's strategy are flipped before being used to update beliefs.
